# The Effect of Node Features on GCN-based Brain Network Classification: An Empirical Study

**DOI:** 10.1101/2022.11.01.514695

**Authors:** Guangyu Wang, Limei Zhang, Lishan Qiao

**Affiliations:** School of Mathematics Science, Liaocheng University, Liaocheng 252000, China; School of Computer Science and Technology, Shandong Jianzhu University, Jinan 250101, China

**Keywords:** Graph convolutional network, Node features, Empirical study, Mild cognitive impairment, Autism spectrum disorder

## Abstract

Brain functional network (BFN) analysis has become a popular technique for identifying neurological/mental diseases. Due to the fact that BFN is a graph, graph convolutional network (GCN) can be naturally used in the classification of BFN. Different from traditional methods that directly use the adjacency matrices of BFNs to train a classifier, GCN requires an additional input — node features. To our best knowledge, however, there is no systematic study to analyze their influence on the performance of GCN-based brain disorder classification. Therefore, in this per, we conduct an empirical study on various node feature measures, including 1) original fMRI signals, 2) one-hot encoding, 3) node statistics, 4) node correlation, and 5) their combination. Experimental results on two benchmark databases show that different node feature inputs to GCN significantly affect the brain disease classification performance, and node correlation usually contributes higher accuracy compared to original signals and manually extracted statistical features.

## I. INTRODUCTION

Resting-state functional magnetic resonance imaging (rs-fMRI) is an effective non-invasive technique for recording spontaneous neural activity in the brain when the subjects are awake and relaxed in the absence of task-specific stimuli. Based on rs-fMRI, the interaction between different regions of interest (ROIs) can be described by brain functional network (BFN) [1] that has become an increasingly important tool to diagnose neurological or mental diseases, such as mild cognitive impairment (MCI) and autism spectrum disorder (ASD) [2].

In consideration of the fact that BFN is a graph, graph convolutional network (GCN) can naturally be used to extract features from BFN for brain disease classification. For example, Parisot *et al*. [3] combined imaging and non-imaging data for population brain analysis using GCN. Arya*et al*. [4] used GCN to fuse structural and functional MRI for ASD classification. Additionally, many improved versions based on GCN were also developed from multi-layer [5, 6] and multi-view [7, 8] to detect brain diseases. Unlike traditional classifiers that are trained directly by the adjacency matrices of BFNs, the GCN-based classification methods need the node feature matrix as an extra input. However, to the best of our knowledge, there is no systematic study to analyze the influence of different node features on the classification results.

In this paper, we use/design four kinds of node feature matrices and evaluate their influence on GCN-based brain disorder classification. The first node feature is original signals (OS) directly extracted from the rs-fMRI. These signals are highly reproducible and provide data sets that can be easily compared across studies. The second node feature is one-hot encoding (OH) that can uniquely identify the location of each node. In addition, we compute several node statistics (NS), including local efficiency, node centrality, and local clustering coefficient, which are concatenated to generate the third type of node features. The fourth node feature is the correlation vector (CV) that reflects the relationship between the current ROI and other ROIs. For a more systematical evaluation, we also discuss the impact of different node feature combinations on the classification results.

Two classification tasks are conducted in this empirical study: 1) MCI identification (identifying subject with MCI from healthy controls) and 2) ASD identification (identifying subject with ASD from healthy controls). Experimental results suggest that different node feature inputs to GCNs have significant effects on brain disease classification performance, and CV usually contributes more than other three features.

The rest of this paper is organized as follows. In Section II, we introduce the relevant concepts and network architecture that will be used in this study. In Section III, we describe the data preparation (including data acquisition and preprocessing) and implementation details. In Section IV, we report the experimental setting and experimental results with a detailed discussion. Finally, we conclude this paper in Section V.

## II. PRELIMINARIES

In this section, we introduce the relevant notation/concepts and architecture of the network used in this study.

### A. Problem Formulation

We represent each subject as a graph *G* = (*V, E*), where *V* = {*v*_1_, *v*_2_, …, *v*_*n*_} denotes the node set, *n* is the number of nodes/ROIs, and *E* = {*e*_1_, *e*_2_, …, *e*_|ε|_} stands for the edge set that is determined by the relationship between different nodes. Accordingly, the node feature matrix and graph adjacency matrix are denoted by *X* ∈ *R* ^*n*×*d*^and *A* ∈ {0, 1}^*n*×*n*^, respectively, where *x*_*i*_ ∈ *R*^*d*^ is the feature of node *v*_*i*_, *d* is the number of feature dimensions and *A*_*ij*_ = 1 if (*v*_*i*_, *v*_*j*_) ∈ *E*, 0 otherwise. We split subjects/graphs into training and test sets. Given an adjacency matrix *A* and a feature matrix *X* in the training sets, we learn a GCN-based classification model and predict the labels of subjects in the test set.

### B. Network Architecture

Our analysis is based on a spectral GCN with two convolution layers [9], which has been empirically verified as an effective architecture for brain disorder identification in several recent works [10, 11, 12]. Its mathematical model can be formulated as follows:

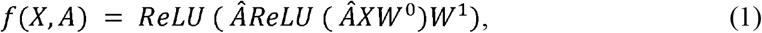

Where 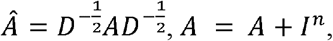 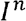 is an identity matrix, and *D* is a diagonal matrix whose diagonal element *D*_*ii*_ = ∑_*j*_ *A*_*ij*_ represents the degree of the *i*-th node, *W* ^0^ and *W* ^1^ are two layers of model parameters that need to be learned from data, and *ReLU* is a nonlinear activation function.

Suppose that *F* = *f* (*X, A*) ∈ *R*^*n*×*d*′^ is the learned representations of nodes, where *f*_*i*_ is the embedding of node *v*_*i*_, *d*′ is the dimension of the feature after embedding. We can further obtain a graph-level representation *H* ^*G*^ = *r*(*F*) ∈ *R*^*d*^′ for by *G* aggregating the node-level embedding. The readout function *r* (·) is a concatenation of the maximum pooling and average pooling operation [13]. It can be formulated as follows:

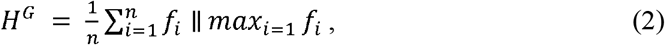

where ‖ denotes concatenation.

## III. EMPIRICAL STUDIES

In this section, we describe the data preparation (including data acquisition and data preprocessing) and implementation details.

### A. Data Preparation

1. *Data acquisition:* In this paper, two publicly available datasets are used to evaluate the effect of node feature on GCN-based method. One is from Alzheimer’s Disease Neuroimaging Initiative (ADNI)^1^ dataset and the other from the Autism Brain Imaging Data Exchange (ABIDE)^2^ dataset. The details of subjects involved in the two datasets are shown in Table 1, including distribution of ASD/MCI, gender (M/F), age and imaging parameters echo time (TE) and repetition time (TR).
2. *Data preprocessing:* All the rs-fMRI data are preprocessed through DPARSF [14] and statistical parametric mapping (SPM12) based on the following standard pipeline: (1) slice timing correction, (2) head motion realignment, (3) nuisance signal regression, including ventricle, white matter signals, and the high-order effect of head motion based on the Friston 24-parameters model, (4) spatial normalization to MNI space, (5) spatial smoothing with the full-width-half-maximum of 4 mm, (6) band-pass temporal filtering from 0.01-0.10Hz. Finally, depending on the AAL atlas, the preprocessed rs-fMRI time series signals are partitioned into 116 ROIs and then averaged respectively to get a representative signal for each ROI.

**TABLE 1.**
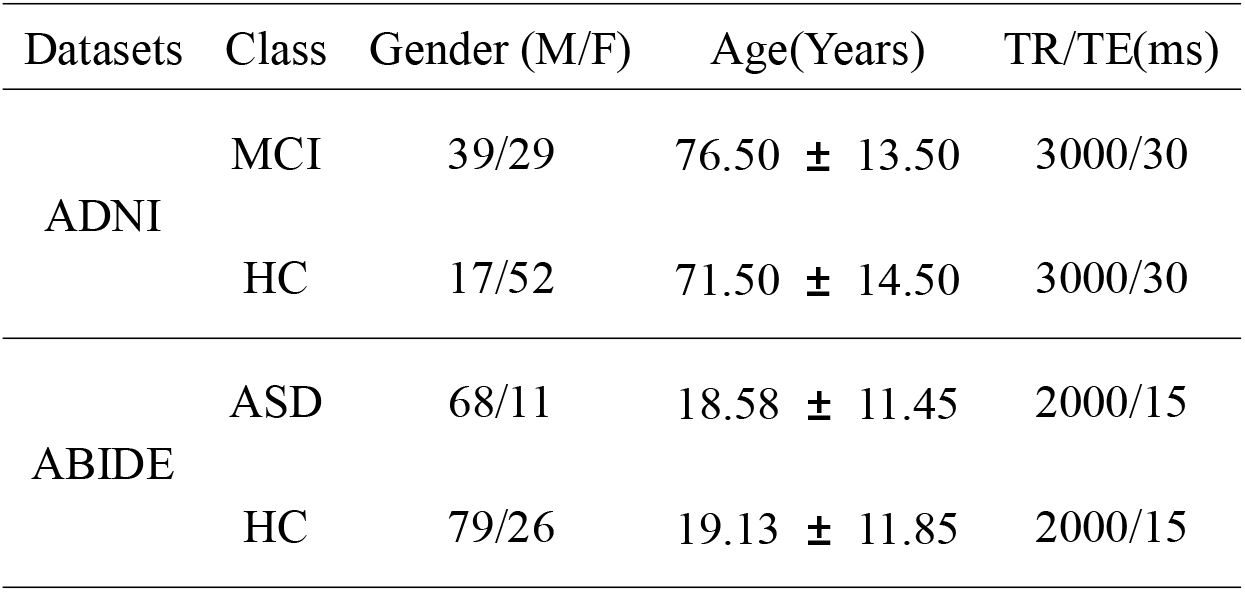
Demographic information of ADNI and ABIDE datasets. M/F: Male/Female. TE/TR: Echo Time/Repetition Time.

### B. Implementation Details

As mentioned earlier, network architecture GCN with two convolution layers, due to its popularity and effectiveness, is used in our study. Furthermore, with the aim of conducting control experiments, we set the embedding dimension as 32 and the number of epochs as 100. Four kinds of node features are involved to verify the classification performance as follows.

1. *Original Signals (OS):* OS capture the spontaneous fluctuations of brain activity associated with different ROIs, which is a manifestation of functional connectivity of the brain.
2. *One-hot Encoding (OH):* We associate each node with a one-hot indicator that uniquely identifies the spatial position of each ROI.
3. *Node Statistics (NS):* We design eight node statistics and concatenate them into a feature vector. In particular, these statistics include three definitions of local clustering coefficients [15], four centralities [16] (*i*.*e*., degree centrality, betweenness centrality, closeness centrality, and eigenvector centrality), as well as local efficiency.
4. *Correlation vector(CV):* CV reflect the relationship between the current ROIs and other ROIs. The most popular measure is Pearson’s correlation (PC) as follows:

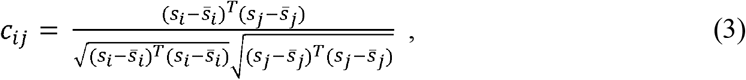

where *s*_*i*_ ∈ *R*^*m*^ (*i* = 1, 2, …, *n*) is the extracted rs-fMRI time series from the *i*-th ROI, *m* is the number of time points, *n* is the number of ROIs, 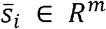 is the mean of *s*_*i*_, and *c*_*ij*_ is the correlation coefficient between the *i*-th and *j*-th ROIs.

### C. Additional Experiments

In general, different BFN estimation methods have a significant influence on the classification performance [17, 18]. Therefore, we discuss the effect of node features based on BFNs estimated by different methods, including PC, sparse representation (SR) [19], and low-rank representation (LR) [20]. The visualizations of three BFNs are shown in Fig.1.

**Fig. 1.**
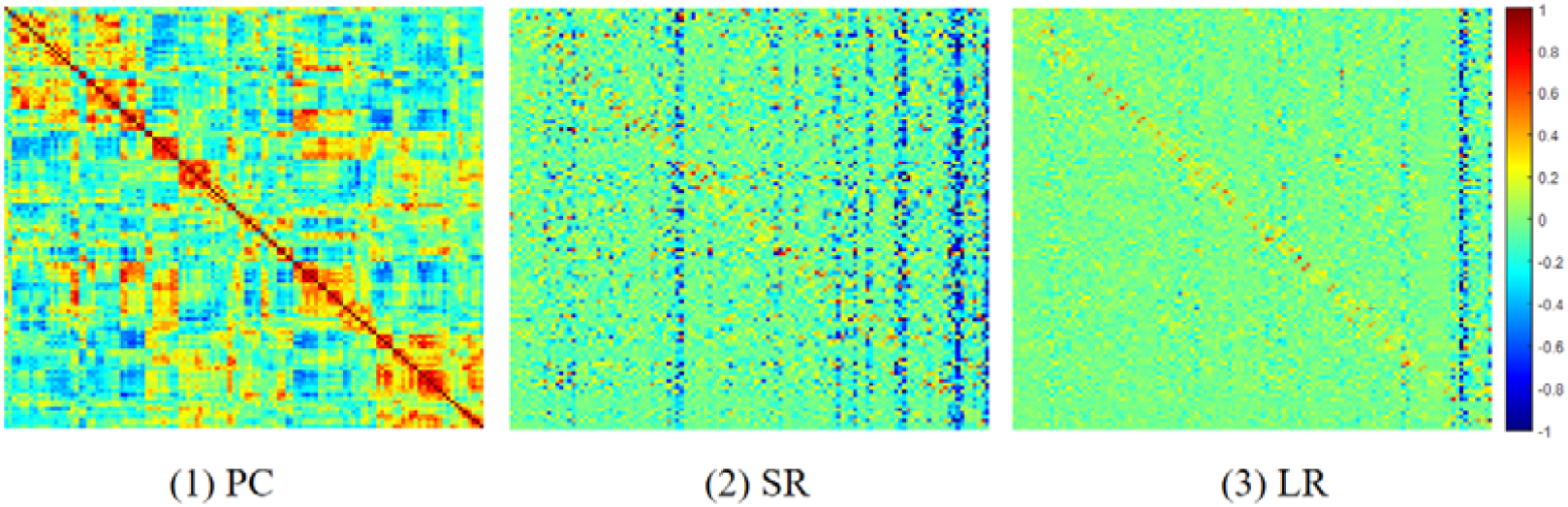
The heat map of adjacency matrices estimated by PC, SR and LR.

## IV. EXPERIMENTS AND RESULTS

### A. Experimental Setting

In this study, we randomly select 80% of the subjects for training and the remaining 20% for testing. For all datasets, we report the mean of 100-run results to evaluate the performance of the involved methods. The performance metrics include

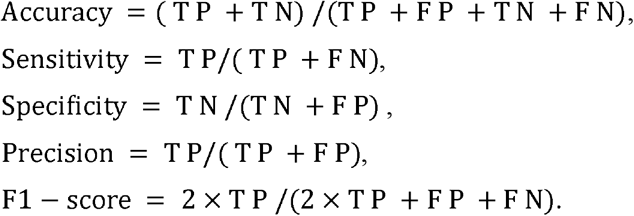

where TP, TN, FP, and FN indicate the true positive, true negative, false positive and false negative, respectively. Additionally, the area under curve (AUC) is adopted for measuring the classification performance.

### B. Results

In Table 2 and Table 3, we report the classification performance corresponding to four kinds of node features and their combinations based on PC adjacency matrix. Then, we show the impact of different node features based on SR and LR adjacency matrices in Fig. 2 and Fig. 3, respectively.

**TABLE 2.**
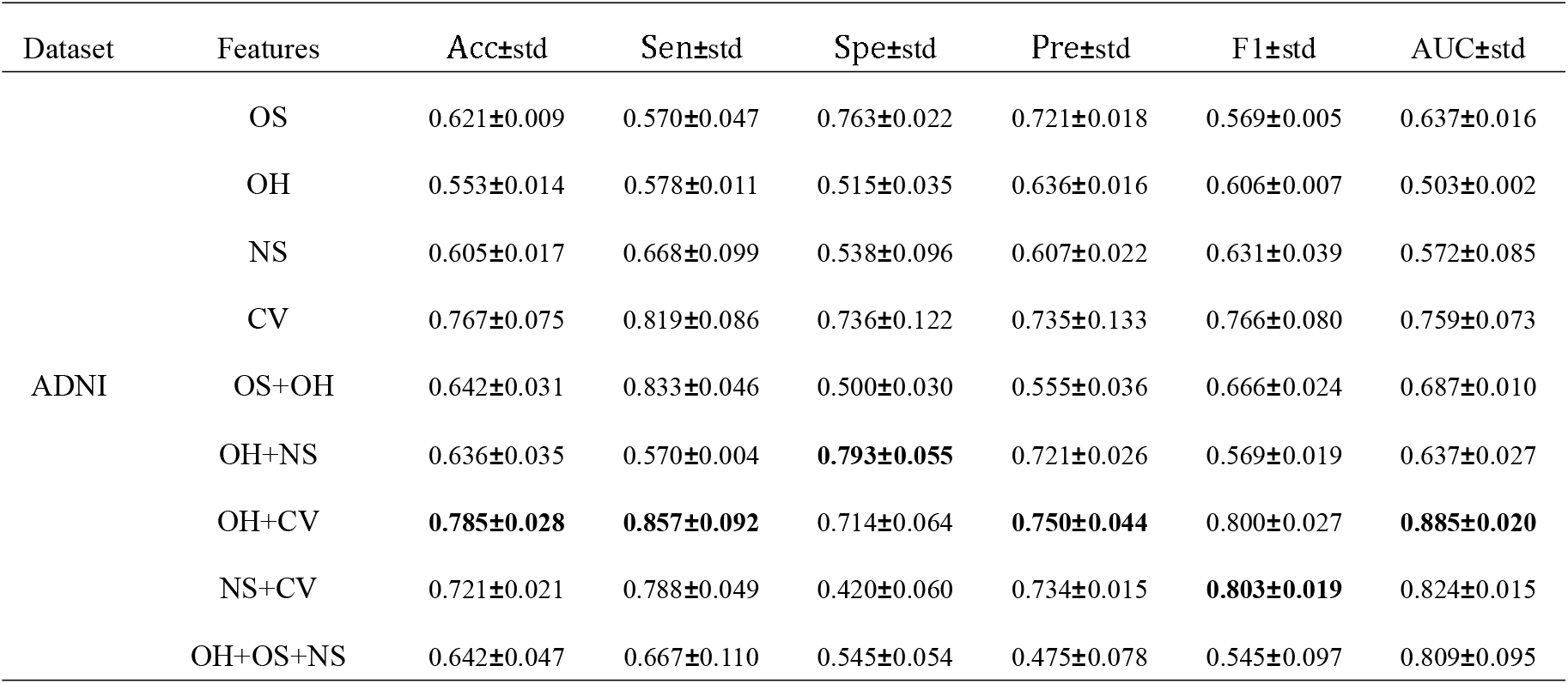
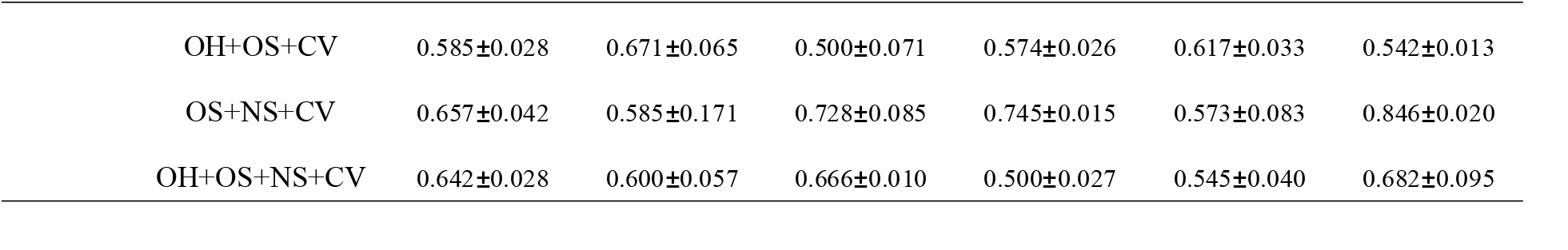
Performance comparison of different node features on both ADNI datasets, with bold values indicating the best results.

| Dataset | Features | Acc±std | Sen±std | Spe±std | Pre±std | F1±std | AUC±std |
| --- | --- | --- | --- | --- | --- | --- | --- |
| ADNI | OS | 0.621±0.009 | 0.570±0.047 | 0.763±0.022 | 0.721±0.018 | 0.569±0.005 | 0.637±0.016 |
| | OH | 0.553±0.014 | 0.578±0.011 | 0.515±0.035 | 0.636±0.016 | 0.606±0.007 | 0.503±0.002 |
| | NS | 0.605±0.017 | 0.668±0.099 | 0.538±0.096 | 0.607±0.022 | 0.631±0.039 | 0.572±0.085 |
| | CV | 0.767±0.075 | 0.819±0.086 | 0.736±0.122 | 0.735±0.133 | 0.766±0.080 | 0.759±0.073 |
| | OS+OH | 0.642±0.031 | 0.833±0.046 | 0.500±0.030 | 0.555±0.036 | 0.666±0.024 | 0.687±0.010 |
| | OH+NS | 0.636±0.035 | 0.570±0.004 | **0.793±0.055** | 0.721±0.026 | 0.569±0.019 | 0.637±0.027 |
| | OH+CV | **0.785±0.028** | **0.857±0.092** | 0.714±0.064 | **0.750±0.044** | 0.800±0.027 | **0.885±0.020** |
| | NS+CV | 0.721±0.021 | 0.788±0.049 | 0.420±0.060 | 0.734±0.015 | **0.803±0.019** | 0.824±0.015 |
| | OH+OS+NS | 0.642±0.047 | 0.667±0.110 | 0.545±0.054 | 0.475±0.078 | 0.545±0.097 | 0.809±0.095 |

|  |  |  |  |  |  |  |
| --- | --- | --- | --- | --- | --- | --- |
| OH+OS+CV | 0.585±0.028 | 0.671±0.065 | 0.500±0.071 | 0.574±0.026 | 0.617±0.033 | 0.542±0.013 |
| OS+NS+CV | 0.657±0.042 | 0.585±0.171 | 0.728±0.085 | 0.745±0.015 | 0.573±0.083 | 0.846±0.020 |
| OH+OS+NS+CV | 0.642±0.028 | 0.600±0.057 | 0.666±0.010 | 0.500±0.027 | 0.545±0.040 | 0.682±0.095 |

**TABLE 3.** Performance comparison of different node features on ABIDE datasets, with bold values indicating the best results.

| Dataset | Features | Acc±std | Sen ± std | Spe ± std | Pre ± std | F1± std | AUC±std |
| --- | --- | --- | --- | --- | --- | --- | --- |
| ABIDE | OS | 0.584±0.079 | 0.619±0.110 | 0.551±0.104 | 0.563±0.101 | 0.589±0.080 | 0.605±0.072 |
|  | OH | 0.538±0.022 | 0.731±0.023 | 0.474±0.026 | 0.574±0.023 | 0.607±0.114 | 0.510±0.095 |
|  | NS | 0.552±0.014 | 0.661±0.087 | 0.538±0.035 | 0.573±0.005 | 0.676±0.040 | 0.536±0.023 |
|  | CV | 0.626±0.055 | 0.662±0.092 | 0.573±0.107 | 0.642±0.012 | 0.653±0.060 | 0.677±0.071 |
|  | OS+OH | 0.705±0.034 | 0.550±0.100 | 0.718±0.081 | **0.701±0.075** | 0.606±0.063 | 0.683±0.010 |
|  | OH+NS | 0.600±0.057 | 0.712±0.148 | 0.450±0.130 | 0.632±0.037 | 0.662±0.085 | 0.589±0.010 |
|  | OH+CV | 0.626±0.015 | 0.500±0.000 | 0.707±0.021 | 0.528±0.014 | 0.560±0.009 | 0.565±0.013 |
|  | NS+CV | **0.747±0.031** | **0.757±0.065** | **0.741±0.069** | 0.636±0.048 | **0.688±0.028** | **0.850±0.009** |
|  | OH+OS+NS | 0.552±0.026 | 0.562±0.037 | 0.690±0.044 | 0.561±0.039 | 0.505±0.035 | 0.568±0.016 |
|  | OH+OS+CV | 0.578±0.019 | 0.525±0.084 | 0.609±0.101 | 0.500±0.018 | 0.500±0.036 | 0.514±0.015 |
|  | OS+NS+CV | 0.600±0.025 | 0.750±0.081 | 0.490±0.044 | 0.518±0.022 | 0.612±0.015 | 0.727±0.075 |
|  | OH+OS+NS+CV | 0.684±0.040 | 0.528±0.110 | 0.733±0.074 | 0.590±0.065 | 0.593±0.097 | 0.782±0.015 |

**Fig. 2.**
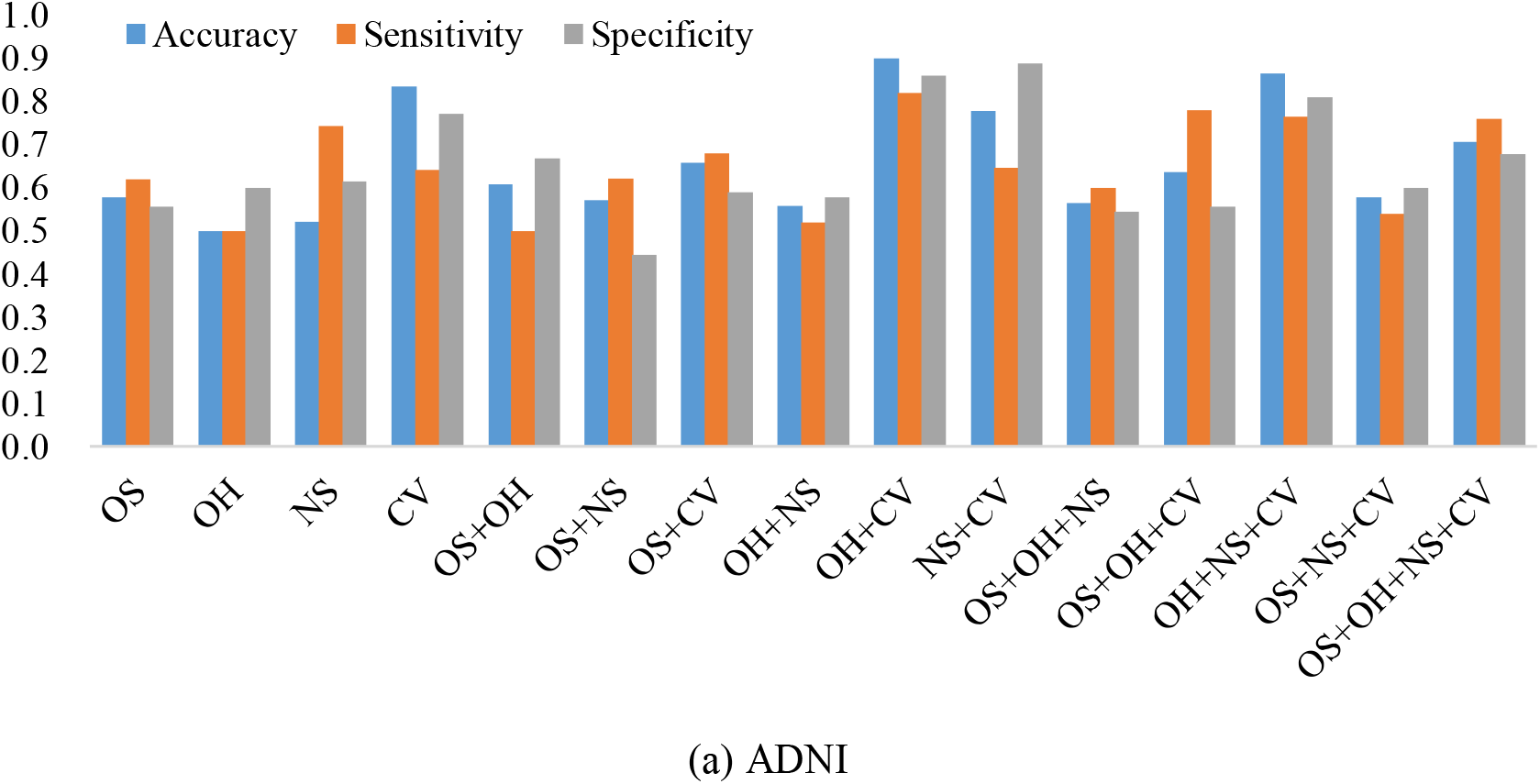
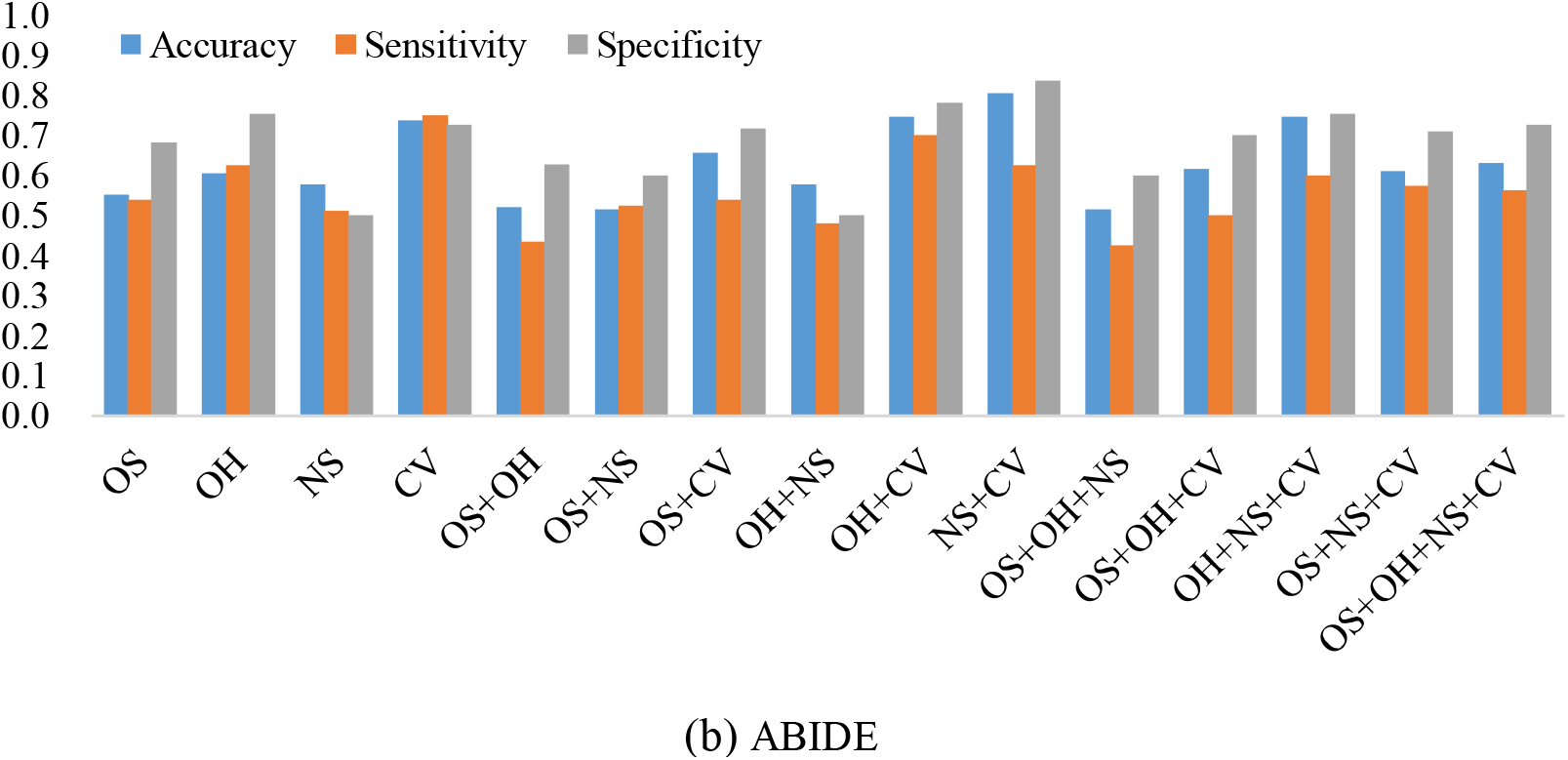
The classification performance corresponding to four node features and their combinations on (a) ADNI and (b) ABIDE datasets, respectively, where the BFNs are estimated by SR.

**Fig. 3.**
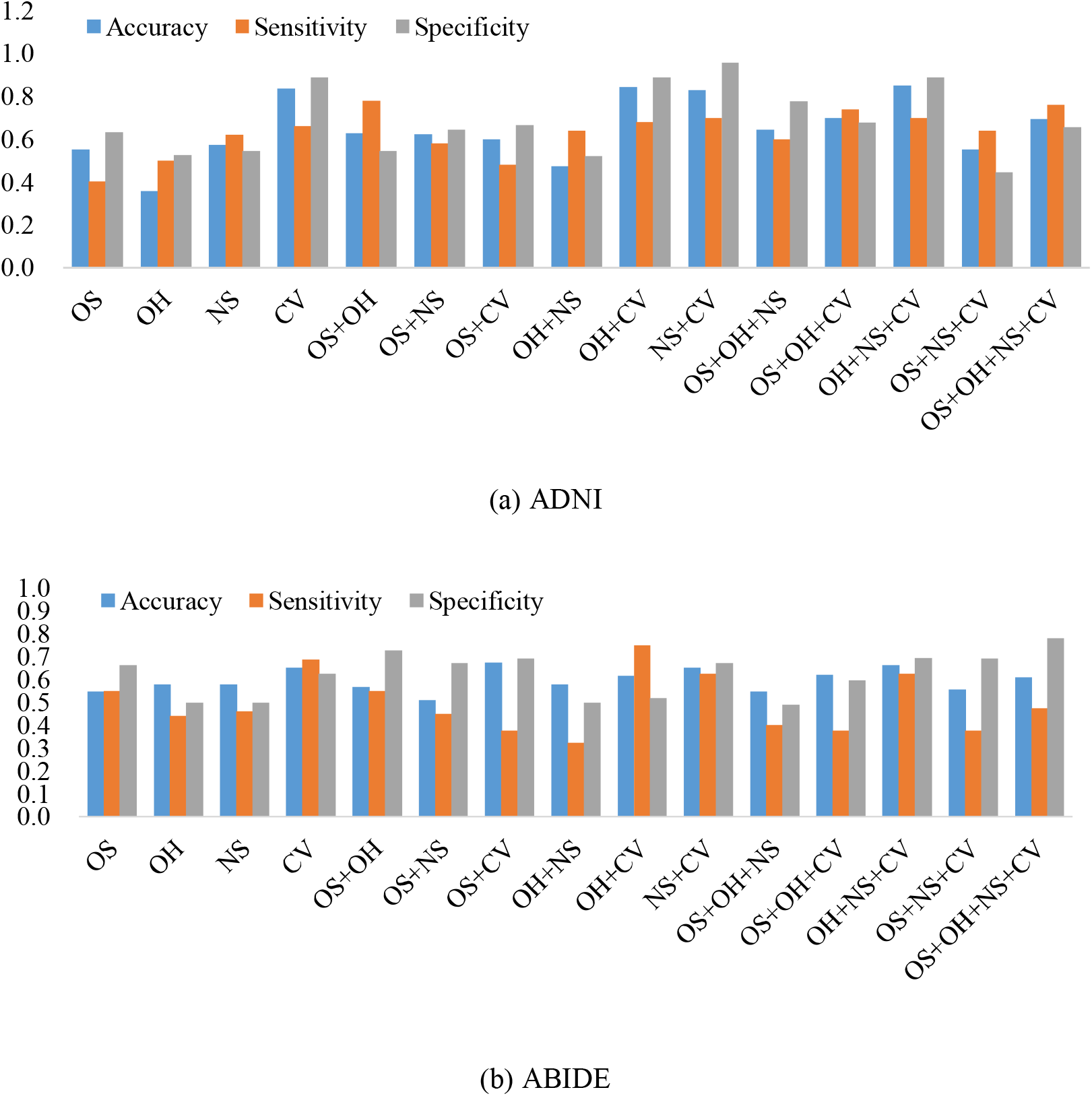
The classification performance corresponding to four node features and their combination on (a) ADNI and (b) ABIDE datasets, respectively, where the BFNs are estimated by LR.

From the above results, we have the following observations.

*First*, for the single node feature, CV can generally result in a higher classification accuracy than the others, and the results are consistent on the two datasets and three differently estimated BFNs (adjacency matrices). This drives us to think that the relationship between ROIs may contain more comparable discriminative information for MCI and ASD identification.

*Second*, the OH gives the worst result among the four single features in most cases, especially on the ADNI dataset, which tells us that only the position encoding of nodes cannot capture informative feature for brain disease classification. Interestingly, however, the combination of OH and CV usually gets the best result. For example, in Table □ and Table □, we note that such a combination achieve the best classification accuracy (78.5% on ANDI dataset and 74.7% on ABIDE dataset, respectively).

*Third*, different feature matrices do have a significant impact on the classification performance. It can also be seen that the designed single features usually cannot achieve the best accuracy. In contrast, when we incrementally concatenate these node features, the accuracy is generally improved. However, concatenating all features cannot achieve the best accuracy, indicating that simply increasing the feature dimension does not guarantee a good classification effect.

*Lastly*, we compare the classification results on different datasets under three adjacency matrices. The results show that on ADNI dataset LR-based adjacency matrix generally leads to a good performance, while on ABIDE dataset SR-based adjacency matrix work better. This suggests that appropriate sparsity is beneficial for brain disease diagnosis compared to the fully connected matrix estimated by PC. This is consistent with many existing studies [21, 22].

## V. CONCLUSION

In this paper, we use/design four kinds of node feature matrices based on rs-fMRI data to empirically evaluate their influence on the GCN-based brain disorder classification tasks. In addition, we conduct control experiments of the feature matrix under three adjacency matrices that correspond to different BFN estimation methods. Experimental results demonstrate that different node features have significant effects on the classification performance. In general, CV, reflecting the relationship between ROIs, or features containing CV tends to result in higher classification performance. Additionally, we also note that combined features usually achieve higher accuracy than single features.

## Footnotes

1 http://adni.loni.usc.edu/

2 http://fcon_1000.projects.nitrc.org/indi/abide

